# Experimental Reptarenavirus Infection of Boa constrictor and Python regius

**DOI:** 10.1101/2020.10.05.327502

**Authors:** U Hetzel, Y Korzyukov, S Keller, L Szirovicza, T Pesch, O Vapalahti, A Kipar, J Hepojoki

## Abstract

Boid inclusion body disease (BIBD) causes losses in captive constrictor snake populations globally. BIBD associates with formation of cytoplasmic inclusion bodies (IB) which mainly comprise reptarenavirus nucleoprotein (NP). In 2017, BIBD was reproduced by cardiac injection of boas and pythons with reptarenaviruses, thus demonstrating a causative link between reptarenavirus infection and the disease. Herein, we report experimental infections of pythons (N=16) and boas (N=16) with three reptarenavirus isolates. First, we used pythons (N=8) to test two virus delivery routes: intraperitoneal injection and tracheal instillation. Independent of the delivery route, we detected viral RNA but no IBs in tissues two weeks post inoculation. Next, we inoculated pythons (N=8) via the trachea. During the four month following the infection snakes showed transient central nervous system (CNS) signs but lacked detectable IB at the time of euthanasia. One of the snakes developed severe CNS signs and we succeeded in re-isolating the virus from the brain of this individual, and could demonstrate viral antigen in neurons. In a third attempt, we tested co-housing, vaccination, and sequential infection with multiple reptarenavirus isolates on boas (N=16). At 10 months post inoculation all except one snake tested positive for viral RNA but none exhibited the characteristic IB. Analysis of the antibody responses demonstrated lower neutralizing but higher anti-reptarenavirus NP titers in experimentally versus naturally reptarenavirus infected boas. Our findings suggest that in addition to reptarenavirus infection, other factors, e.g. the antibody response, contribute to BIBD pathogenesis.

**IMPORTANCE:** A 2017 study demonstrated cardiac reptarenavirus injection to induce boid inclusion body disease (BIBD) in pythons and boas. In the present study, we experimentally infected pythons and boas with reptarenavirus via either intraperitoneal injection or tracheal instillation. We found both virus delivery routes to result in infection; though the latter could reflect the natural route of infection. In the experimentally infected snakes, we did not find evidence of inclusion body (IB) formation, characteristic to BIBD, in pythons or in boas. Most of the snakes (11/12) studied were reptarenavirus infected after ten-month follow up, which suggests that they could eventually have developed BIBD. We further found differences between the antibody responses of experimentally and naturally reptarenavirus infected snakes, which could indicate that the pathogenesis of BIBD involves factors additional to reptarenavirus infection. As snakes are poikilotherm, also the housing conditions could have an effect.

## INTRODUCTION

There are descriptions of a plague called boid inclusion body disease (BIBD) in captive snake populations since the 1970s (1). The disease mostly affects members of the families *Boidae* and *Pythonidae*, and may lead to eradication of entire snake collections (1, 2). BIBD manifests itself in a variety of clinical conditions, such as neurological signs including regurgitation, head tremors, loss of coordination, and as abnormal skin shedding, secondary bacterial infections, and neoplastic diseases (1, 2). The pathognomonic hallmark of BIBD is the formation of cytoplasmic ultrastructurally electron-dense and histologically eosinophilic inclusion bodies (IBs) in almost all cell types (3, 4). The standard ante mortem diagnosis of BIBD relies on the IB detection in blood smears or tissue biopsies (1, 5, 6). Even before the causative agent of BIBD was identified, the IBs were found to consist mainly of a 68 kDa protein, unknown at the time (4). In 2012/2013 the findings of three independent groups linked BIBD with arenavirus infection (7-9). Furthermore, we and others could demonstrate that the “68 kDa protein” actually represents the arenavirus nucleoprotein (NP) (5, 7, 9).

The identification of arenaviruses in snakes led to establishment of two new genera, *Mammarenavirus* (previously known arenaviruses) and *Reptarenavirus* (BIBD-associated arenaviruses), within the family *Arenaviridae* (10). We and others then made the observation that snakes with BIBD most often, if not always, carry several reptarenavirus L and S segments (11, 12). These studies dramatically expanded the number of fully sequenced reptarenavirus L segments, from four to approximately 150 (11, 12). Currently, the L segment of close to 30 reptarenavirus species are known based on the ICTV’s (International Committee on Taxonomy of Viruses) species demarcation criteria (species share <76% nt identity) (10). The high genetic diversity makes nucleic acid-based diagnostic approaches to BIBD challenging, and thus the detection of reptarenavirus antigen (nucleoprotein, NP) serves as an alternative (6). Additionally, increasing evidence indicates that reptarenavirus infection is not always associated with detectable IBs (6, 13, 14), suggesting that the BIBD pathogenesis may involve additional factors. For example, we demonstrated vertical transmission of co-infecting reptarenavirus L and S segments with concurrent presence of IBs (15), and thus congenital, peri- or neonatal infection could be a prerequisite for IB formation. We also identified Haartman Institute Snake virus-1 (HISV-1) in a snake with BIBD (12), which led to establishment of the third arenavirus genus, *Hartmanivirus* (16, 17). Later, we observed that snakes with BIBD fairly often also carry hartmaniviruses, however, we could so far not link hartmanivirus infection to BIBD (14, 18, 19).

At present, the family *Arenaviridae* comprises four genera: *Mammarenavirus, Reptarenavirus, Hartmanivirus*, and *Antennavirus* (17). The genome of all except antennaviruses is a bisegmented negative-sense RNA (20). The L segment of mammarenaviruses and reptarenaviruses encodes an RNA-dependent RNA polymerase (RdRp) and a zinc finger matrix Z protein (ZP), while the S segment encodes the glycoprotein precursor (GPC) and NP (21). The L segment of hartmaniviruses lacks the ORF for ZP (18).

The literature describes at least three attempts to reproduce BIBD *in vivo*. In 1994 Schumacher and co-workers injected two 3-month-old Burmese pythons (*Python molurus bivitattus*) with cell-free supernatants of cultured primary kidney cells of a *Boa constrictor* with BIBD (3). Both developed CNS signs, leading to the death of the first animal at six weeks post inoculation, and euthanasia of the second after 10 weeks (3). The pathological examination revealed a non-suppurative, lymphocyte dominated encephalitis with neuronal degeneration in both animals. IBs were only found in the second animal, and only in neurons in the brain and in the pituitary gland, but not in other organs. The authors’ attempts to re-isolate and identify the causative agent were unsuccessful (3). In 2000 Wozniak and co-workers infected four *B. constrictors* intraperitoneally with filtered liver homogenate from a BIBD positive donor and observed IBs in hepatocytes 10 weeks post infection (4). They succeeded in isolating the IBs and in generating a monoclonal antibody against the “68 kDa protein” (most likely reptarenavirus NP, in retrospect), but could not characterize the causative agent (4). At the time of the Schumacher and Wozniak studies BIBD was suspected to be caused by an unknown retrovirus. In 2017, Stenglein and co-workers reported to have reproduced BIBD in *Python regius* and *B. constrictor* by cardiac injection of purified reptarenavirus (22). The authors diagnosed classical BIBD, as defined by IB formation, in boas but did not observe IBs in pythons (22). Furthermore, while the boas remained clinically healthy for two years after infection, the pythons developed severe CNS signs within two months (22). These findings highlight the complexity of BIBD pathogenesis, and provide further evidence that the disease outcome might vary not only between viruses but also between snake species.

Herein we report the results of a series of experimental infections of pythons (*P. regius*) and boas (*B. constrictor*). When we initiated the experimental infections, in 2013, our primary aim was to demonstrate the etiologic relationship between reptarenavirus infection and BIBD. We tested two different routes, intracoelomic and tracheal, for inoculation of the snakes with purified cell culture-grown reptarenaviruses. We also studied the possibility of vaccinating the snakes against reptarenavirus infection, and potential transmission during co-housing. During the third set of experimental infections, we learned that snakes with BIBD are often co-infected with several reptarenavirus species (12), and decided to attempt inoculating snakes with multiple reptarenaviruses in both co- and superinfection setups. We subjected all snakes to a full post mortem examination, used RT-PCR to detect viral RNA, immunohistology (anti-reptarenavirus NP) to detect viral antigen, ELISA for detecting anti-reptarenavirus antibodies in snakes, and vesicular stomatitis viruses (VSV) pseudotyped with reptarenavirus glycoproteins to detect neutralizing antibodies (NAbs) in the snakes.

## MATERIALS AND METHODS

### Ethics statement

The experimental infection was approved by the National Animal Experiment Board (Eläinkoelautakunta, ELLA) of Finland (permit number, ESAVI/4690/04.10.07/2013). All animals were euthanized according to Schedule 1 procedures to minimize suffering.

### Cells, viruses and purification of viruses

The continuous *B. constrictor* kidney cell line, I/1Ki generated and maintained as described (9, 23), served for virus production and virus re-isolation attempts from tissues and blood of the experimentally infected animals. One isolate used in this study, University of Helsinki virus (UHV), was initially described in (9), but we later found it to actually comprise two reptarenaviruses, UHV-1 (GenBank accessions: KR870020.1 and KR870011.1) and Aurora borealis virus-1, ABV-1 (GenBank accessions: KR870021.1 and KR870010.1), at roughly equal amounts as judged by reads obtained by NGS (12). The other isolate (T10404) used was from snake no. 5 in (9), later named University of Giessen virus-1 (UGV-1; GenBank accession numbers: KR870022.1 and KR870012.1). The propagation, purification, and storage of UHV and UGV-1 preparations used for inoculation have been described in (23). Re-isolations of virus from infected snakes were done by overlaying I/1Ki cells (80-90% confluent) with EDTA blood and tissue homogenates of brain, lung, liver, kidney, and heart for 24 h at 30 °C, followed by media exchange and 10-14 d incubation at 30 °C. The cells were analyzed for viral antigen expression by western blotting. The cell culture supernatant collected at 5 and 10 days post infection (dpi) was cleared by centrifugation (3000 x g, 5 min), 0.45 μm filtered, and the viruses pelleted by ultracentrifugation (27.000 x g, 5 °C, 2 h) through a 1 ml 30% sucrose cushion (in phosphate-buffered saline, PBS, pH 7.4) in a SW41 rotor (Beckman coulter). The pelleted virus material was re-solubilized in PBS and analyzed by sodium docecyl sulphate-polyacrylamide gel electrophoresis (SDS-PAGE) and western blotting. Virus titration was done as described for hantaviruses (24).

### Animals and infection

BIBD negative animals in blood smear (16 *Python regius* obtained were obtained from a commercial German breeder and 16 *Boa constrictor* from a private Swiss breeder). The animals were housed in aerated plastic boxes (Smartstore Classic 15, Orthex Group, dimensions 40×30×19 cm), held in temperature (between 27-30 °C) and day-light (12 h of light) controlled ERHET cabins. The humidity (approximately 60-80%) inside housing boxes was maintained by evaporation from a water supply.

For the initial trial (see Table 1), involving eight juvenile (2 months of age) pythons (*P. regius*) from a single clutch, three snakes were infected with UHV preparation (contains UHV-1 and ABV-1, but for simplicity referred to as UHV), three with UGV-1, and two remained as controls. The infected animals in both groups were inoculated as follows: one received 5,000 fluorescent focus forming units (fffus) intracoelomic, one 50,000 fffus intracoelomic, and one 50,000 fffus instilled into the trachea (the volume of inoculum was 500 μl in PBS). The control animals received 500 μl PBS intracoelomic and intratracheal respectively. The snakes were monitored daily for clinical signs, and euthanized at 14 dpi.

**Table 1.** First experimental infection.

| **Animal** | **Species** | **Age** | **Inoculation route** | **Virus and dose** | **RT-PCR (Brain, Lung, Blood)** | **Blood smear** | **Histology** | **IHC** |
| --- | --- | --- | --- | --- | --- | --- | --- | --- |
| 1.1 | *P. regius* | Juvenile (~2 Mo) | Trachea and coelomic cavity | mock (PBS) | Neg, Neg, Neg | Neg | Neg | Neg |
| 1.2 | *P. regius* | Juvenile (~2 Mo) | Trachea and coelomic cavity | mock (PBS) | Neg, Neg, Neg | Neg | Neg | Neg |
| 1.3 | *P. regius* | Juvenile (~2 Mo) | coelomic cavity | UGV-1, 5000 fffus | UGV-1, Neg, Neg | Neg | Neg | Neg |
| 1.4 | *P. regius* | Juvenile (~2 Mo) | coelomic cavity | UGV-1, 50000 fffus | UGV-1, Neg, Neg | Neg | Neg | Neg |
| 1.5 | *P. regius* | Juvenile (~2 Mo) | Trachea | UGV-1, 50000 fffus | UGV-1, UGV-1, Neg | Neg | Neg | Neg |
| 1.6 | *P. regius* | Juvenile (~2 Mo) | coelomic cavity | UHV (UHV-1 and ABV-1), 5000 fffus | Neg, Neg, Neg | Neg | Neg | Neg |
| 1.7 | *P. regius* | Juvenile (~2 Mo) | coelomic cavity | UHV (UHV-1 and ABV-1), 50000 fffus | Neg, Neg, Neg | Neg | Neg | Neg |
| 1.8 | *P. regius* | Juvenile (~2 Mo) | Trachea | UHV (UHV-1 and ABV-1), 50000 fffus | Neg, UHV-1 and ABV-1, ABV-1 | Neg | Neg | Neg |

In the second experimental infection (see Table 2), again involving eight 2-month-old pythons (*P. regius*) from a single clutch obtained from a commercial German breeder, four pythons (*P. regius*) received the UHV preparation, two received UGV-1, and two were administered the equivalent amount of PBS. Virus inocula (50,000 fffus for both UGV-1 and UHV inoculations, all diluted in 0.5 ml of PBS) were instilled into the trachea. The snakes were monitored daily and fed at one to two week intervals. At 44 dpi, two juvenile (4 months of age) boas (*B. constrictor*) were included into the experiment, one was co-housed with a UHV (animal 2.4) and the other with a UGV-1 (animal 2.8) inoculated python. All snakes were monitored daily for any clinical signs. The animals were euthanized as follows: animal 2.1 at 118 dpi, 2.2, 2.3 and 2.7 at 22 dpi, 2.5 at 30 dpi (due to severe CNS signs), 2.4 at 69 dpi, 2.6 and 2.8 at 117 dpi. The two co-housed boas (2.9 and 2.10) were also euthanized at 117 dpi (73 days post initiation of co-housing), at the scheduled end of the experiment.

**Table 2.**
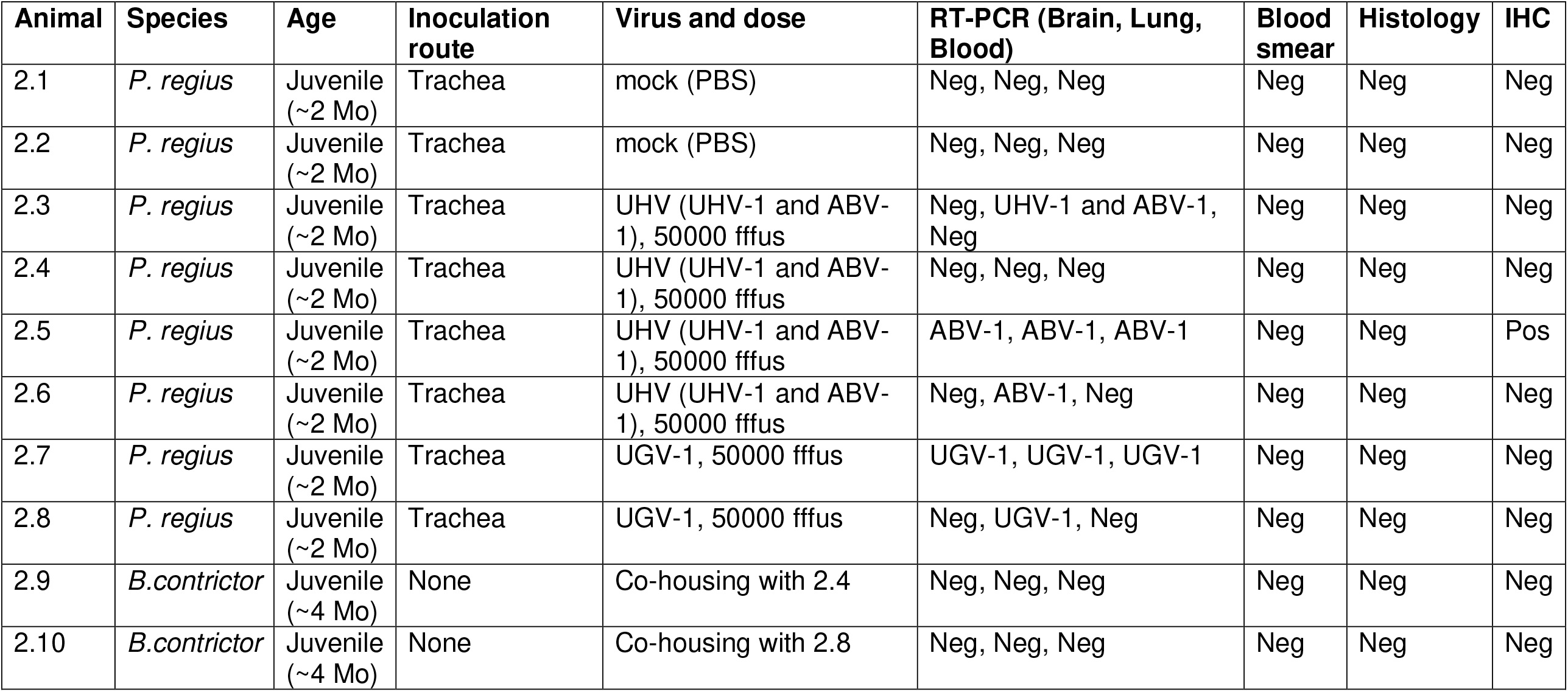
Second experimental infection.

For the third experimental infection (see Table 3), we received a clutch of 16 *B. constrictors*, of which three were immunized with purified UHV inactivated by addition of Triton X-100 (to a final concentration of 0.2% v/v; animals 3.4, 3.5 and 3.6), and one (animal 3.3) with recombinant UHV NP (described in (23)). Briefly, at day 0 the animals were subcutaneously administered either approximately 10,000,000 fffus of detergent-inactivated UHV or 0.1 mg of recombinant UHV NP emulsified in Freund’s incomplete adjuvant (ThermoFisher Scientific), the total volume per individual was 125 μl. At 13 and 26 d after the initial administration, boosters with a similar dose were administered. At 74 days post initial immunizations eight boas (animals 3.3-3.10, including the vaccinated ones, 3.3-3.6) received 250,000 fffus of UHV, two boas (animals 3.11 and 3.12) received 125,000 fffus of both UHV and UGV-1, and two boas (animals 3.13 and 3.14) received 250,000 fffus of UGV-1, by tracheal instillation. At 116 dpi, two vaccinated snakes were administered 250,000 fffus of UGV-1 (animals 3.3 and 3.4), and the two snakes initially inoculated with UGV-1 (animals 3.13 and 3.14) were placed into boxes with UHV-inoculated snakes for co-housing (animals 3.9 and 3.10, respectively). The snakes were monitored daily for any clinical signs and fed at one to three week intervals.

**Table 3.**
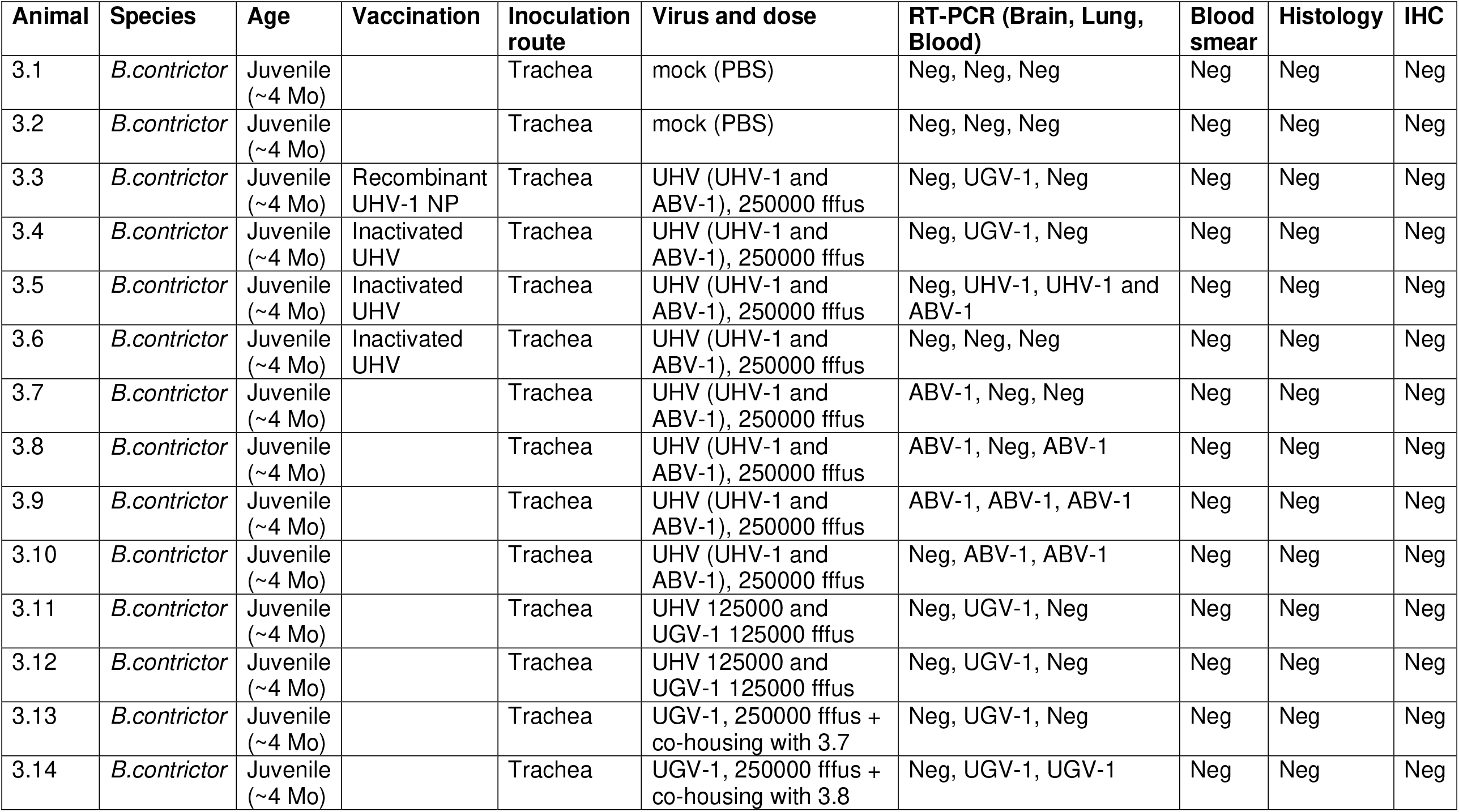
Third experimental infection.

Prior to virus inoculation, a blood sample had been collected from the tail vein of each animal. Snakes were euthanized by decapitation after sedation by exposure to CO_2_. A blood sample was collected and animals were necropsied and organ samples collected immediately into Trizol (Life Technologies, for RT-PCR) and into paraformaldehyde (PFA; 4% solution in PBS, for histology and immunohistology), and were fresh frozen at −70 °C for virus isolation and further analyses.

### SDS-PAGE and immunoblotting

SDS-PAGE and immunoblotting were done as described (9, 23). The antibodies against UHV nucleoprotein (NP), described in (23), were used for the detection in immunoblottings. The visualization of immunoblots probed (at 1:10,000 dilution) with goat anti-rabbit IR800Dye (LI-COR biosciences) or goat anti-rabbit AlexaFluor 680 (Invitrogen) was done using the Odyssey Infrared Imaging System (LI-COR bioscience).

### Reverse transcription-polymerase chain reaction (RT-PCR) and Sanger sequencing

RNA isolation from tissue and blood samples was done as described (15). The RT-PCRs for UHV-1, ABV-1, and UGV-1 L and/or S segments were done initially using primers and protocol described in (15), and the RT-PCR products were analysed by standard agarose gel electrophoresis visualized by GelRed Nucleic Acid Stain (Biotium) and subjected to Sanger sequencing (core facility of the Haartman Institute, University of Helsinki, Finland). The isolated RNAs were later re-analyzed using a one step Taqman assay with the following primers and probes targeting the S segment: UGV-1 probe 6-Fam-CTCGACAAGCGTGGGCGGAGG-BHQ-1, UGV-1-fwd 5’-CAAGAAAAACCACACTGCACA-3’, UGV-rev 5’-AACCTGTTGTGTTCAGTAGT-3’, UHV-1 probe 6-Fam-TCCTCTGCCGCAAAAGACTATGTCACAG-BHQ-1, UHV-1-fwd 5’ -ACAAACTGAATAAGACTGCTGCATT-3’, UHV-1-rev 5’-AGGGCTATACACACATAGTTGGATG-3’, ABV probe 6-Fam-CATGAATTCTTCATCGACATCAGAAACCG-BHQ-1, ABV-1-fwd 5’-CCGTACTGCACAACTGATGATG-3’, ABV-1-rev 5’-AGCAACACAGGAGTAACCTGTCAC-3’, and following the TaqMan Fast Virus 1-Step Master Mix (ThermoScientific) product guidelines.

### Histology and immunohistochemistry (IHC)

For histology and IHC, samples of brain, lung, liver, kidney, pancreas, spleen, small intestine, and heart were fixed in PFA for 48 h and routinely paraffin wax embedded. Sections (3-4 µm) were prepared and stained with hematoxylin-eosin (HE). For all RT-PCR-positive animals, consecutive sections were prepared and subjected to IHC for viral NP, employing the recently described broadly cross-reactive rabbit anti-pan-reptarenavirus antiserum (25), following a previously described protocol (9, 23).

### Focus reduction neutralization test (FRNT) using replication incompetent vesicular stomatitis virus (VSV) pseudotyped with reptarenavirus GPs

Production of single cycle replication, GP deficient, recombinant VSV expressing the enhanced green fluorescent protein (scrVSVΔG-eGFP) pseudotyped with different reptarenavirus GPs was done as described (26). Each pseudotyped scrVSVΔG-eGFP batch was titrated with a 10-fold dilution series on a 96-well plate of clean I/1Ki cells, and the dilution yielding 50-150 fluorescent cells was selected for FRNT. To demonstrate neutralizing antibodies (NAbs) against UHV-1, UGV-1, and ABV-1, EDTA plasma was prepared from the blood samples (14) and incubated with the pseudotyped VSVs at 30 °C for 60 min prior to laying the virus-serum mixture onto 80-90% confluent I/1Ki cells. After 2 h incubation at 30 °C, the virus-plasma mixture was replaced with fresh complemented medium, and the plate was incubated for 16-24 h at 30 °C. Infected cells were enumerated using fluorescence microscopy. All experiments were performed in triplicate. Plasma samples of 24 naturally reptarenavirus infected snakes from an earlier study (14) were analysed using VSVs pseudotyped with S5-like, Tavallinen suomalainen mies virus-2 (TSMV-2), and UGV-1 GPs. These reptarenavirus S segments had been found in the collection, and the snakes had been analysed by RT-PCR for their presence at the time of sampling (14). The neutralizing titer was determined as the plasma dilution that induced at least a 50% reduction in the number of fluorescent foci.

## RESULTS

### Selection of the infection route

In the first experimental infection involving eight juvenile, approximately 2-month-old *P. regius*, we tested whether the route of admission would affect the course of infection. Cell culture adaptation is for many viruses known to cause virus attenuation. Thus we decided to use two virus preparations, UHV (containing UHV-1 and ABV-1) which has been propagated in tissue cultures for >8 years, and UGV-1, after a single passage. The viruses were purified by ultracentrifugation, diluted in PBS and used to inoculate each three snakes (Table 1), two into the coelomic cavity (5,000 and 50,000 fffus), the third via the respiratory route, by instillation into the trachea (50,000 fffus), to best mimic a possible natural route of infection. Some animals exhibited slight lethargy, but none showed clinical signs during the following two weeks (Fig. 1). At the time of euthanasia, two weeks post inoculation, RT-PCR confirmed UGV-1 infection of the brain regardless of the route of infection, whereas only tracheal instillation of UHV resulted in detection of viral RNA, and only in the lung, at the end of the experiment (Table 1). None of the snakes showed IB formation in blood or tissues, and there was no evidence of viral NP expression including the brain.

**Figure 1.**
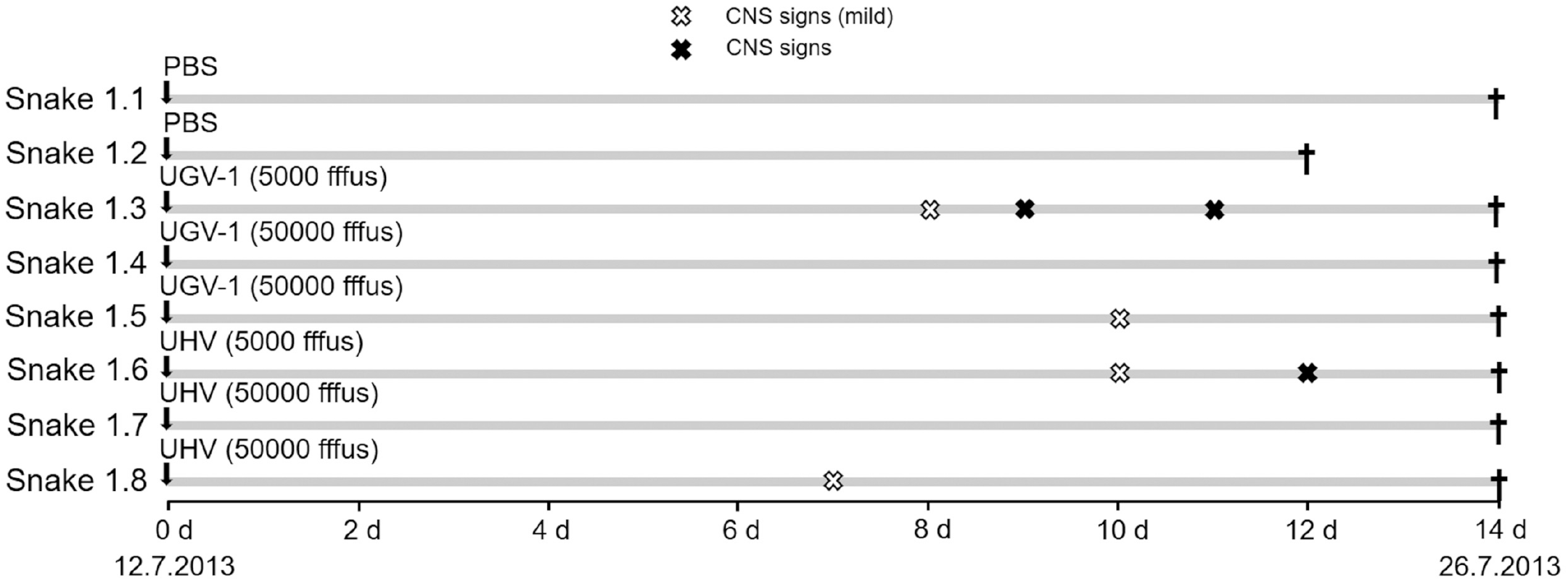
A schematic representation of the first experimental infection timeline. The experiment included eight pythons (*P. regius*), which were monitored for 14 days post inoculation. The vertical arrows indicate inoculation. White (mild tremor) and black (tremor) X-marks in the infection timeline mark the observed CNS signs. The black crosses indicate euthanasia.

### Experimental infection of a group of pythons (*P. regius)*

After demonstrating inoculation via the trachea to be effective in the initial trial, we decided to employ tracheal inoculation in the subsequent experiments because it likely reflects the natural route of infection. For the experiment, we inoculated four juvenile pythons (animals 2.3 to 2.6) at the age of approximately 2 mo with UHV, two (2.7, 2.8) with UGV-1, and two control animals (2.1 and 2.2) with PBS (Table 2, Fig. 2). We monitored the snakes daily for signs of disease, and at 19 dpi animals 2.3, 2.6, 2.7, and 2.8 showed mild head tremor (Fig. 2). Animal 2.6 also exhibited unphysiological tail postures. At 22 dpi two of these snakes (2.3 and 2.7) were euthanized as scheduled, together with one control snake (2.2). Both animals 2.3 and 2.7 were found to be infected; animal 2.3 was RT-PCR positive for both inoculated viruses (UHV-1, ABV-1), but only in the lung. It did not show IB formation in any tissue (Table 2). Animal 2.6 exhibited viral RNA in brain and lungs, but neither IB formation nor viral antigen expression.

**Figure 2.**
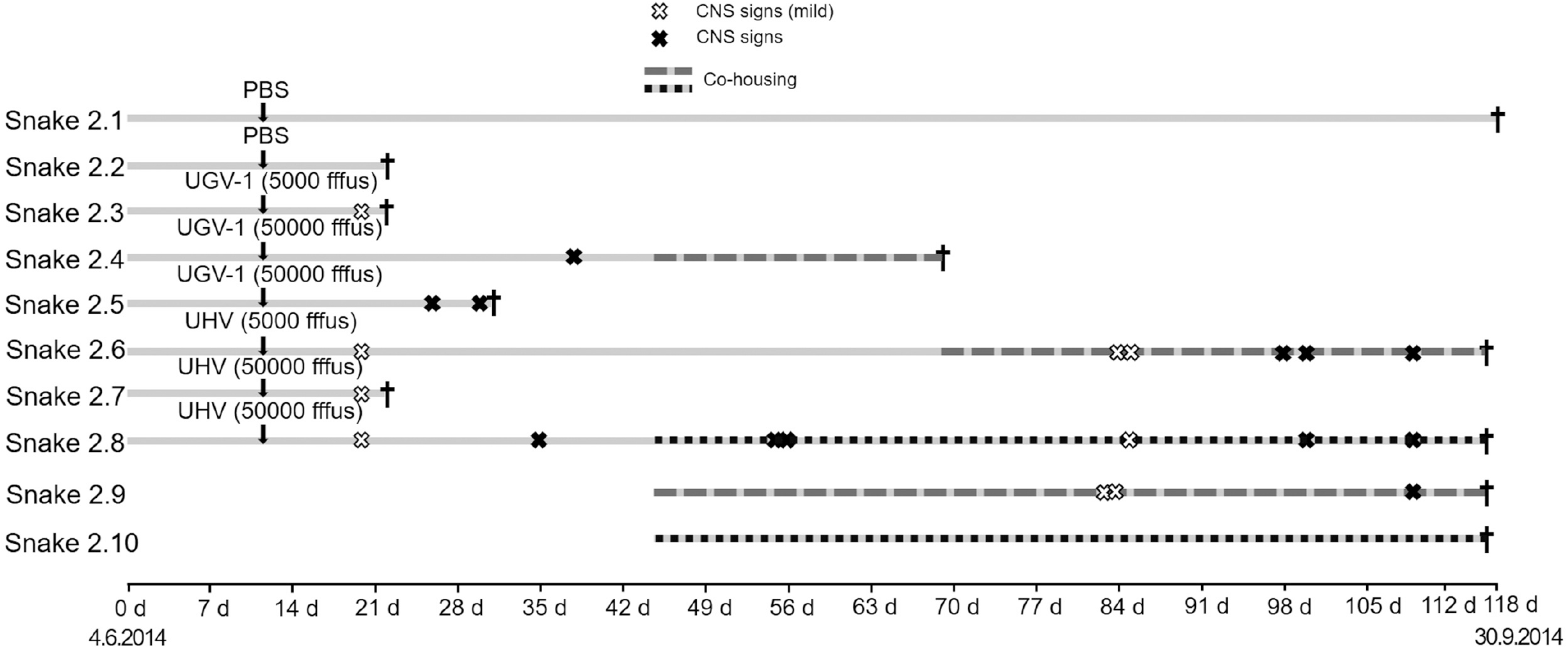
A schematic representation of the second experimental infection timeline. The experiment included eight pythons (*P. regius*) and two boas (*B. constrictor*), which were monitored up to 118 days post inoculation. The vertical arrows indicate inoculation. The observed CNS signs are marked by white (mild tremor) and black (tremor) X-marks, and the co-housing of snakes by shading of the infection timeline. The black crosses indicate euthanasia.

At 25 dpi we observed neurological signs in animal 2.5 (body balance and coordination problems). At 29 dpi, during feeding, animal 2.5 showed tremor and lethargy, and had severe difficulties to swallow its feed (a frozen mouse); the snake had to be euthanized the following day, since the clinical signs had worsened (Fig. 2). We found virus by RT-PCR in both lungs and brain and could purify the virus from I/1Ki cells inoculated with a brain homogenate. We could not detect IBs in blood cells, but the animal exhibited reptarenavirus NP expression in neurons in the brain and in cells with the morphology of macrophages and/or dendritic cells in spleen and thymus (Fig. 3, Table 2). At 34 dpi, animal 2.8 showed CNS signs (body balance and coordination problems) and at 37 dpi animal 2.4 showed head tremors; in both snakes, the clinical signs improved and vanished during the following days (Fig. 2). At 43 dpi, having received 16 juvenile *B. constrictor* snakes, we decided to investigate whether co-housing with experimentally infected *P. regius* would result in virus transmission across the two species. We placed one boa each in the box of one python (animal 2.9 to animal 2.4, and animal 2.10 to animal 2.8). At 54 dpi animal 2.8 again showed CNS signs (tremors and disorientation). The next day animal 2.4 also showed similar signs. However, the clinical signs of animal 2.8 improved during the following days (Fig. 2). At 61 dpi animal 2.6 which had shown mild CNS signs early after inoculation (day 19), had diarrhea, but was otherwise in good condition. At 69 dpi we sacrificed animal 2.4 since the mild CNS signs had by then persisted for two weeks. The animal did not exhibit IBs or viral antigen expression, and we did not find reptarenavirus RNA in the tissues studied. The boa (animal 2.9) that had been co-housed with this animal was moved to the box of animal 2.6.

**Figure 3.**
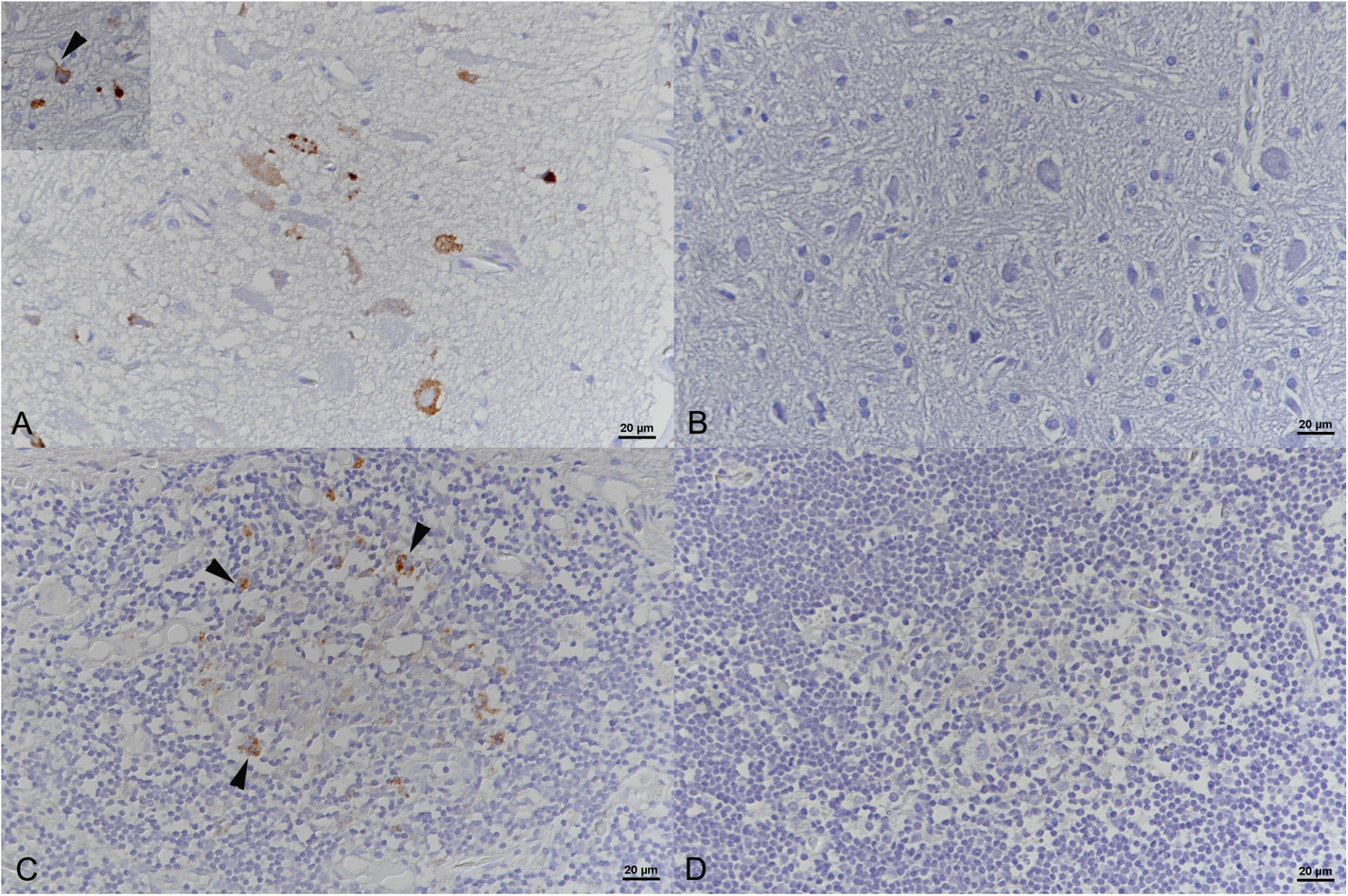
Brain and spleen, *Python regius*, euthanized 22 days post intratracheal instillation of UHV (animal 2.5). Immunohistochemistry (IHC) shows for reptarenavirus nucleoprotein in the cytoplasm of neurons (arrowhead) in the brain (**A**) and in macrophages/dendritic cells (arrowheads) in the spleen (**C**). Negative control slides (**B** and **D**). IHC employing a broadly cross-reactive rabbit anti-pan-reptarenavirus antiserum (25), hematoxylin counterstain. Bars = 20 µm.

At 83 to 85 dpi animal 2.6 showed lethargy and the co-housed boa (animal 2.9) displayed abnormal tail postures. In addition, animal 2.8 was lethargic at 85 dpi. The clinical signs of all three snakes improved during the following days, but from 98 dpi onwards, animal 2.6 was again lethargic. At 100 dpi, animal 2.8 showed similar lethargy, but again, both snakes improved during the following days. At 109 dpi, when feeding, both again showed CNS signs (mild tremor) and refused to feed. The boa (animal 2.9) co-housed with a python (animal 2.6) showed similar signs and difficulties in eating. At 117 dpi we decided to euthanize these three animals (2.6, 2.8, 2.9) as well as the boa (animal 2.10) that had shared the box with the python (animal 2.8). At 118 dpi we sacrificed the remaining control animal, snake 2.1. All except the control animal (2.1) and the co-housed boas were found reptarenavirus positive by RT-PCR in the lung, and animal 2.7 carried the virus also in the brain. The histological analysis did not reveal IB formation in any of these animals; reptarenavirus antigen expression was also not detected (Table 2).

### Vaccination, and experimental infection challenge of *B. constrictors*

Since the first two rounds of experimental infections had been unsuccessful in terms of replicating IB formation, we finally decided to attempt inoculation of boas (*B. constrictor*), since the virus isolates originate from this species. We also decided to attempt vaccination prior to inoculations and used purified UHV lysate (three animals: 3.3 to 3.5) or recombinant UHV-1 NP ((23), snake 3.6). We gave the first vaccinations the day after the boas had arrived (−74 dpi), i.e. the start day for the python-boa co-housing experiment. Around two (−61 dpi) and four weeks (−48 dpi) later we boosted animals 3.3 to 3.6 with the same antigens (Table 3, Fig. 4).

**Figure 4.**
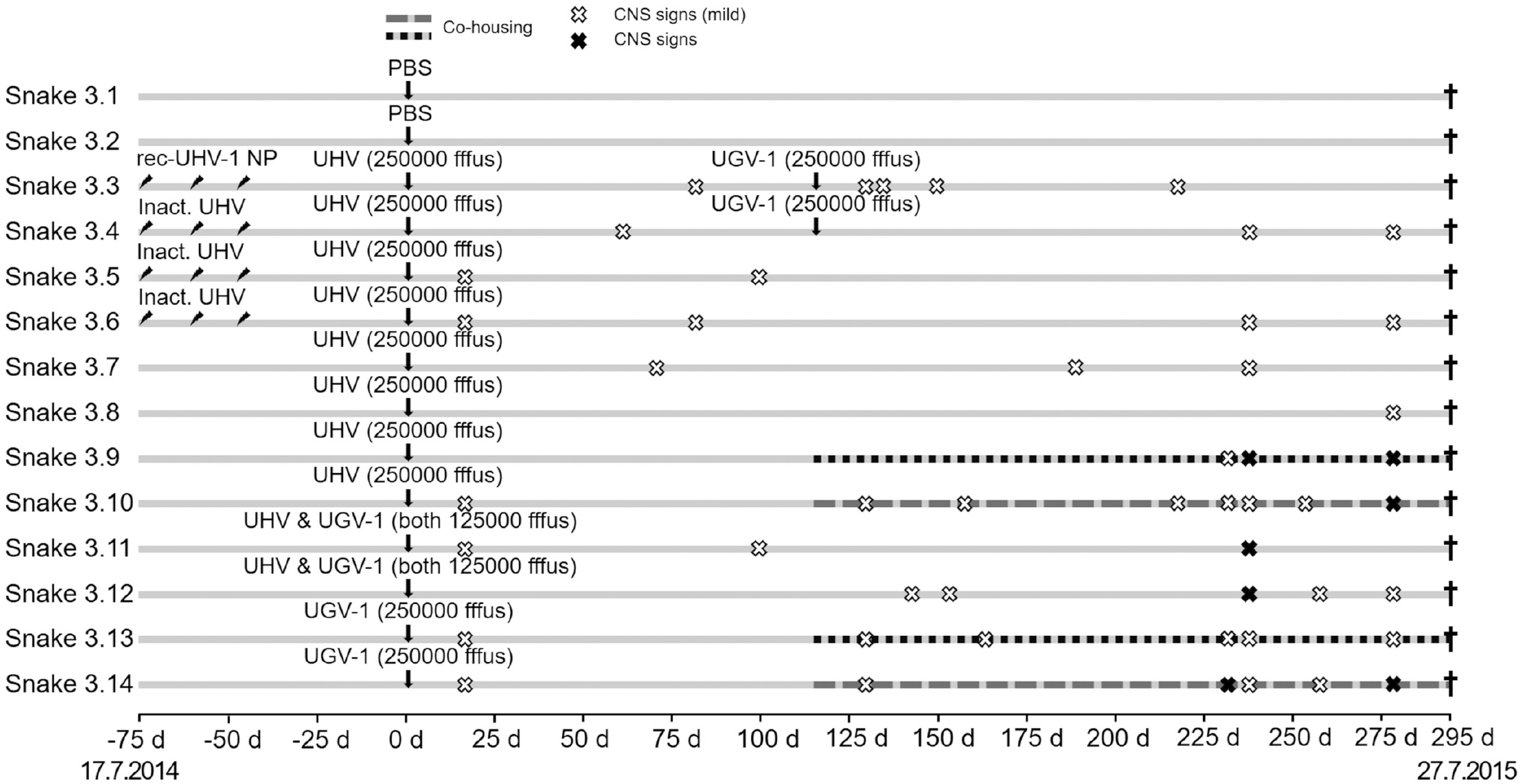
A schematic representation of the third experimental infection timeline. The experiment included 14 boas (*B. constrictor*), which were monitored up to 295 days (∼10 months) post inoculation. The inclined arrows indicate immunization, and the vertical arrows indicate inoculation time points. White (mild tremor) and black (tremor) X-marks indicate the observed CNS signs, and the co-housing of snakes by shading of the infection timeline. The black crosses indicate euthanasia.

Since we had successfully infected the pythons but been unsuccessful inducing IB formation, we decided to increase the amount of input virus and chose to use an infectious dose that was five-fold higher than that used earlier. We also wanted to attempt co-infection of some snakes with UHV and UGV-1, but were at this point unaware that our UHV preparation was indeed a mix of ABV-1 and UHV-1. We inoculated the vaccinated boas (animals 3.3 to 3.6) and four non-vaccinated boas (animals 3.7 to 3.10) with UHV (250,000 fffus/snake), two boas (animals 3.11 and 3.12) with a mix of UHV and UGV-1 (125,000 fffus/snake of each), and two boas (animals 3.13 and 3.14) with UGV-1 (250.000 fffus/snake), Table 3. At 14 dpi we observed mild tremors in animals 3.5, 3.6, 3.10, 3.11, 3.13, and 3.14, but the signs waned the following days. Afterwards, mild CNS signs were observed occasionally; at 57 dpi (animal 3.4, stargazing), at 59 dpi (animal 3.4, tremor), at 68 dpi (anima 3.7, tremor), and at 79 dpi (animals 3.3 and 3.6, tremor), Fig. 4.

After discovery that snakes with BIBD often carry several reptarenavirus L and S segments (12), we decided to super-infect some snakes by re-inoculation at 115 dpi: animals 3.3 and 3.4 (originally inoculated with UHV) received 250,000 fffus/snake of UGV-1. At this point, we had also detected using next-generation sequencing that our UHV preparation actually contains two viruses (ABV-1 and UHV-1) and decided to retry virus transmission during co-housing. Therefore, at 115 dpi, we placed animal 3.9 (UHV inoculated) in the box of animal 3.13 (UGV-1 inoculated), and animal 3.10 (UHV inoculated) in the box of animal 3.14 (UGV-1 inoculated). We continued monitoring the snakes and observed the following intermittent clinical signs that affected all animals at some point: at 127 dpi (animals 3.3, 3.10, 3.13, 3.14, mild tremors), at 132 dpi (3.3, mild tremor), at 140 dpi (3.12, mild tremor and disorientation), at 149 dpi (3.3, disorientation), at 152 dpi (3.12, stargazing), at 155 dpi (3.10, tremors and disorientation), at 161 dpi (3.13, mild tremor), at 186 dpi (3.7, disorientation), at 215 dpi (3.10, mild tremor; 3.3 disorientation), at 229 dpi (3.9, 3.10, 3.13, and 3.14, mild tremors; 3.14 lethargy), at 235 dpi (3.4, 3.6, 3.7, 3.9 to 3.11, 3.13, and 3.14, mild tremor; 3.9, 3.11 and 3.12, lethargy), at 251 dpi (3.10 and 3.11, mild tremors; 3.4, 3.8, 3.10 and 3.12, lethargy), at 261 dpi (3.12, tremor), at 265 dpi (3.12, tremor; 3.14, stargazing), and at 276 dpi (3.4, 3.6, 3.8 to 3.10 and 3.12 to 3.14, lethargy), Fig. 4.

At 294 dpi we euthanized animals 3.3 to 3.8, and at 295 dpi we euthanized 3.9 to 3.14, as scheduled. Using RT-PCR we could confirm that all animals except snake 3.4 were infected. They carried viral RNA in one or more of the tissues studied; five animals (3.4, 3.8, 3.10 to 3.12) were also found to be viremic. None of the animals exhibited IB or reptarenavirus antigen in any tissue or the blood, Table 3.

### Immune response against reptarenavirus NP in experimentally and naturally infected snakes

Unlike Stenglein and colleagues (22), we did not succeed in inducing IB formation, the hallmark of BIBD (1, 3-7, 9), by experimental reptarenaviruses infection in python or boas. We recently learned that reptarenavirus infected snakes with IBs in blood cells have lower levels of anti-reptarenavirus antibodies as compared to reptarenavirus infected snakes without IBs (14). Thus we compared the antibody responses of the experimentally infected snakes to responses in naturally reptarenavirus infected boas (the latter using a panel of 24 plasma samples available from an earlier study (14)) using tools developed earlier (14, 23, 26, 27). We used ELISA with purified UGV-1 as the antigen to determine the level of IgM and IgY antibodies against reptarenavirus NP (Table 4). None of the python sera produced signal in the ELISA, most likely indicating lack of cross-reactivity of our anti-boa immunoglobulin reagents to python immunoglobulins, rather than lack of antibodies. The ELISA results show that the uninfected boas that served as control snakes did not have anti-reptarenavirus NP antibodies, suggesting that the snakes had not been in contact with reptarenaviruses prior to vaccination and/or inoculation (Table 4 and Fig. 5A). The result also indicates that the vaccinations with both inactivated UHV preparation and recombinant UHV-1 NP had induced the formation of anti-NP antibodies (Table 4 and Fig. 5A-B). We took the serum samples from the vaccinated snakes at day 74 after the initial vaccination, which is the likely reason for the presence of IgY but not IgM class anti-NP antibodies. At the end of the experiment, approximately 10 months post virus challenge, all snakes had IgY class anti-NP antibodies (Table 4 and Fig. 5B). Their anti-NP IgY levels were slightly higher than those of the naturally infected snakes of which some were also anti-NP IgM positive (Tables 4 and 5A). We then compared the results of snakes without IB to those of snakes with IB (i.e. confirmed BIBD) and observed that the latter had lower levels of anti-NP antibodies (Table 4 and Fig. 5A-B), a finding reported earlier (14).

**Table 4.** Antibody responses in experimentally as compared to naturally reptarenavirus infected boas.

| Animal | Anti-NP IgG (ELISA) | Anti-NP IgM (ELISA) | Blood smear (0-3) | UGV-1 neut. titer | UHV-1 neut. titer | ABV-1 neut. titer | TSMV-2 neut. titer | S5-like neut. titer | RT-PCR |
| --- | --- | --- | --- | --- | --- | --- | --- | --- | --- |
| 3.1 0-bleed | 0,151 | 0,052 | 0 | 0 | 0 | 0 | ND | ND | Neg |
| 3.1. final | 0,201 | 0,058 | 0 | 0 | 0 | 0 | ND | ND | Neg |
| 3.2 0-bleed | 0,199 | 0,067 | 0 | 0 | 0 | 0 | ND | ND | Neg |
| 3.2. final | 0,109 | 0,026 | 0 | 0 | 0 | 0 | ND | ND | Neg |
| 3.3. post. imm. | 3,227 | 0,069 | 0 | ND | ND | ND | ND | ND | Neg |
| 3.3 final | 3,311 | 0,109 | 0 | 250 | 150 | 75 | ND | ND | UGV-1 |
| 3.4. post. imm. | 2,838 | 0,059 | 0 | ND | ND | ND | ND | ND | Neg |
| 3.4 final | 3,092 | 0,109 | 0 | 150 | 100 | 75 | ND | ND | UGV-1 |
| 3.5. post. imm. | 1,303 | 0,080 | 0 | ND | ND | ND | ND | ND | Neg |
| 3.5 final | 1,901 | 0,045 | 0 | 500 | 500 | 200 | ND | ND | UHV-1, ABV-1 |
| 3.6. post. imm. | 3,212 | 0,026 | 0 | ND | ND | ND | ND | ND | Neg |
| 3.6 final | 3,103 | 0,041 | 0 | 200 | 400 | 50 | ND | ND | Neg |
| 3.7 final | 3,018 | 0,081 | 0 | 200 | 150 | 400 | ND | ND | ABV-1 |
| 3.8 final | 3,177 | 0,066 | 0 | 400 | 0 | 800 | ND | ND | ABV-1 |
| 3.9 final | 3,177 | 0,042 | 0 | 400 | 200 | 800 | ND | ND | ABV-1 |
| 3.10 final | 2,910 | 0,039 | 0 | 400 | 200 | 50 | ND | ND | ABV-1 |
| 3.11 final | 3,267 | 0,059 | 0 | 400 | 50 | 0 | ND | ND | UGV-1 |
| 3.12 final | 3,319 | 0,060 | 0 | 400 | 400 | 400 | ND | ND | UGV-1 |
| 3.13 final | 3,372 | 0,262 | 0 | 400 | 50 | 400 | ND | ND | UGV-1 |
| 3.14 final | 3,370 | 0,049 | 0 | 400 | 1600 | 200 | ND | ND | UGV-1 |
| Nat. inf. 1 | 3,042 | 0,061 | 0 | 200 |  |  | 0 | 6400 | UGV, TSMV-2 |
| Nat. inf. 2 | 1,839 | 0,085 | 0 | 1000 |  |  | 50 | 3200 | TSMV-2 |
| Nat. inf. 3 | 1,186 | 0,047 | 0 | 500 |  |  | 400 | 3400 | S5, TSMV-2 |
| Nat. inf. 4 | 0,045 | 0,008 | 0 | 700 |  |  | 6400 | 6400 | Neg |
| Nat. inf. 5 | 0,150 | 0,089 | 2 | 90 |  |  | 100 | 800 | UGV, S5, TSMV-2 |
| Nat. inf. 6 | 0,621 | 0,045 | 0 | 800 |  |  | 350 | 1600 | S5 |
| Nat. inf. 7 | 0,222 | 0,116 | 2 | 450 |  |  | 50 | 350 | UGV, S5 |
| Nat. inf. 8 | 1,914 | 0,026 | 0 | 750 |  |  | 400 | 3500 | Neg |
| Nat. inf. 9 | 0,161 | 0,021 | 1 | 400 |  |  | 300 | 1600 | UGV, S5 |
| Nat. inf. 10 | 2,348 | 0,122 | 0 | 400 |  |  | 0 | 2200 | S5 |
| Nat. inf. 11 | 2,270 | 0,257 | 0 | 600 |  |  | 800 | 750 | Neg |
| Nat. inf. 12 | 0,054 | 0,018 | 3 | 800 |  |  | 1700 | 2400 | UGV, S5 |
| Nat. inf. 13 | 1,546 | 0,101 | 0 | 200 |  |  | 0 | 300 | UGV, TSMV-2 |
| Nat. inf. 14 | 0,085 | 0,012 | 2 | 1700 |  |  | 50 | 1200 | UGV, S5, TSMV-2 |
| Nat. inf. 15 | 1,828 | 0,262 | 1 | 1700 |  |  | 300 | 1200 | UGV |
| Nat. inf. 16 | 1,588 | 0,132 | 0 | 800 |  |  | 75 | 3700 | UGV, S5, TSMV-2 |
| Nat. inf. 17 | 1,049 | 0,416 | 3 | 1600 |  |  | 0 | 6400 | UGV, S5, TSMV-2 |
| Nat. inf. 18 | 0,699 | 0,015 | 1 | 300 |  |  | 250 | 6000 | UGV, S5, TSMV-2 |
| Nat. inf. 19 | 0,217 | 0,017 | 0 | 250 |  |  | 250 | 2800 | S5, TSMV-2 |
| Nat. inf. 20 | 0,034 | 0,060 | 0 | 200 |  |  | 200 | 300 | S5, TSMV-2 |
| Nat. inf. 21 | 1,581 | 0,078 | 0 | 400 |  |  | 400 | 450 | S5, TSMV-2 |
| Nat. inf. 22 | 0,020 | 0,045 | 1 | 400 |  |  | 250 | 1700 | UGV, S5, TSMV-2 |
| Nat. inf. 23 | 1,813 | 0,054 | 0 | 200 |  |  | 200 | 1200 | UGV, S5, TSMV-2 |
| Nat. inf. 24 | 0,983 | 0,706 | 1 | 75 |  |  | 100 | 1700 | S5, TSMV-2 |

Blood smear: 0-3 refers to the size of the IBs, 0=no inclusions and 3=large and/or numerous IBs.

**Figure 5.**
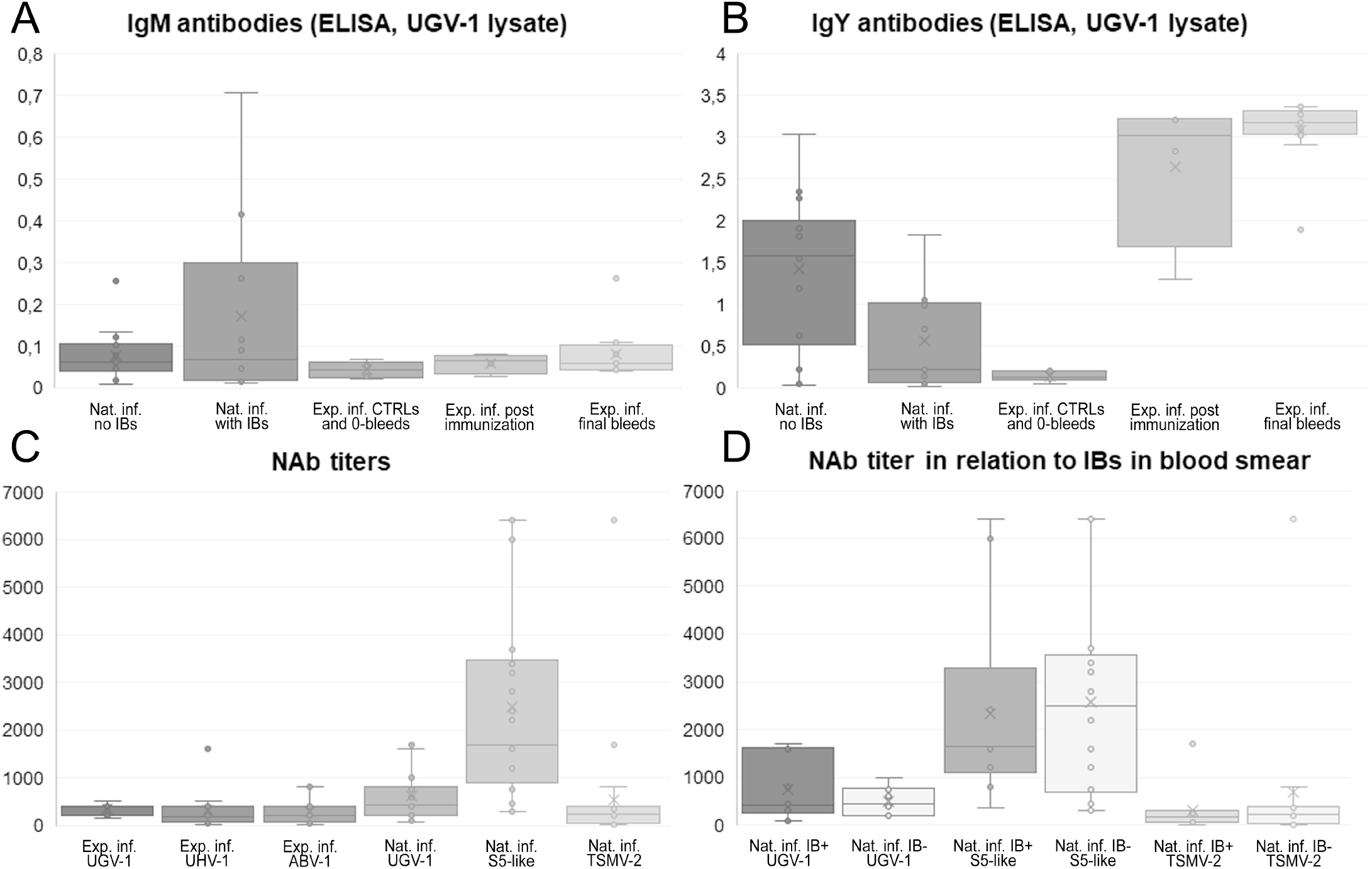
Antibody responses in experimentally versus naturally infected boas (*B. constrictor*). **A)** A box plot of IgM class antibodies against reptarenavirus NP (using concentrated UGV-1 lysate as the antigen). The boxes from left represent: naturally infected snakes without IBs, naturally infected snakes with IBs, the 0-bleeds collected prior to immunization or inoculation, samples collected following immunization, and samples collected from the experimentally infected snakes at the time of euthanasia. The y-axis represents OD_450 nm_ values as the ELISA readout. **B)** A box plot of IgY class antibodies against reptarenavirus NP (using concentrated UGV-1 lysate as the antigen). The boxes from left represent: naturally infected snakes without IBs, naturally infected snakes with IBs, the 0-bleeds collected prior to immunization or inoculation, samples collected following immunization, and samples collected from the experimentally infected snakes at the time of euthanasia. The y-axis represents OD_450 nm_ values as the ELISA readout. **C)** A box plot of neutralizing antibody (NAb) titers as studied using VSV pseudotypes with reptarenavirus glycoproteins. The boxes from left represent: neutralizing antibodies against UGV-1 in experimentally infected snakes, neutralizing antibodies against UHV-1 in experimentally infected snakes, neutralizing antibodies against ABV-1 in experimentally infected snakes, neutralizing antibodies against UGV-1 in naturally infected snakes, neutralizing antibodies against S5-like glycoproteins in naturally infected snakes, and neutralizing antibodies against TSMV-2 in naturally infected snakes. The y-axis represents the last dilution producing 50% reduction in the number of fluorescent foci. **D)** A box plot of neutralizing antibody (NAb) titers as studied using VSV pseudotypes with reptarenavirus glycoproteins in naturally infected snakes with and without IBs. The boxes from left represent: neutralizing antibodies against UGV-1 in snakes with IBs, neutralizing antibodies against UGV-1 in snakes without IBs, neutralizing antibodies against S5-like glycoproteins in snakes with IBs, neutralizing antibodies against S5-like glycoproteins in snakes without IBs, neutralizing antibodies against TSMV-2 in snakes with IBs, and neutralizing antibodies against TSMV-2 in snakes without IBs. The y-axis represents the last dilution producing 50% reduction in the number of fluorescent foci.

### Reptarenavirus NAbs in experimentally and naturally infected snakes

As the analysis of anti-reptarenavirus NP antibodies indicated potential differences in the immune response of experimentally versus naturally infected snakes, we wanted to compare the NAb response in both groups of snakes. Using a fluorescent replication-defective recombinant vesicular stomatitis virus (rVSV-ΔG*eGFP) system, we had previously generated single-round infectious particles pseudotyped with reptarenaviral GPCs (ABV-1, UHV-1, UGV-1, S5-like, and TSMV-2) (26) which we employed to determine the 50% focus reduction neutralization titer (FRNT50) for the sera. We studied the boa sera from the third experimental infection against ABV-1, UHV-1 and UGV-1 GP bearing pseudotypes, and used UGV-1, S5-like, and TSMV-2 GP bearing pseudotypes for the sera of the naturally infected snakes from our previous study (14). The latter were selected based on the result of RT-PCRs targeting the respective S segments (14). The results indicate differences in the neutralizing titers of the experimentally infected snakes (Table 4 and Fig. 5C-D). Four snakes (animals 3.3, 3.4, 3.10 and 3.11) showed the highest FRNT50 titer against UGV-1, two (animals 3.6 and 3.14) against UHV-1, and three (animals 3.7, 3.8 and 3.9) against ABV-1. Snake 3.5 showed equal FRNT50 titers for UGV-1 and UHV-1, snake 3.12 for all studied viruses, and snake 3.13 for UGV-1 and ABV-1. Two of the snakes, 3.8 (inoculated with UHV) and 3.10 (inoculated with both UHV and UGV-1), did not show NAbs against UHV-1 and ABV-1, respectively, but 3.9 had a good NAb response against UGV-1. Animal 3.12 had the highest recorded neutralizing titer against UHV-1, reaching a value of 1,600. Two snakes, animals 3.1 and 3.2, mounted the weakest NAb response with the highest titer reaching 250, while for other experimentally infected snakes the highest titers were at or above 400.

The naturally infected snakes showed much higher NAb titers, for most animals the highest titer was clearly above 1,000 and for some we recorded neutralizing titers as high as 6,400 (Table 4 and Fig. 5C). We further observed that a high neutralizing titer against a given virus did not provide neutralization against other reptarenaviruses (see e.g. animals #5, #23, #27, and #37 in Table 4), which suggests that the level of cross-neutralization might be low.

## DISCUSSION

BIBD has remained an enigmatic disease for decades. Before identifying reptarenaviruses as the likely causative agent, at least two studies had reproduced the disease under experimental conditions using cell culture isolated causative agent (3, 4). In the 1994 report by Schumacher and coauthors, one of the two infected Burmese pythons (*P. bivitattus*) developed severe CNS signs and died six weeks post inoculation, the second was euthanized ten weeks post inoculation due to severe CNS signs (3). The authors did observe IBs in the brain of one of the snakes, however, they failed to re-isolate the infectious agent (3). Retrospectively, it is possible that the re-isolation *per se* was successful but the authors merely failed to detect the causative agent since reptarenaviruses do not induce a cytopathic effect in cell culture. In the second study, by Wozniak and colleagues, the authors inoculated four juvenile boas (*B. constrictor*) with a liver homogenate from a snake with BIBD and included two equal-sized control groups: non-inoculated, and inoculated with a liver homogenate from a healthy boa (4). By ten weeks post inoculation, all four snakes inoculated with liver homogenate from a BIBD positive snake had developed IBs in the liver, but none showed clinical signs of BIBD during the one-year surveillance period (4). After identification of reptarenaviruses as the most likely etiological agent for BIBD, Stenglein and colleagues performed an experimental infection on two pythons (*P. regius*) and two boas (*B. constrictor*) (22). The pythons developed CNS signs within two months post inoculation, but did not show IB formation and only the brain samples tested positive for the viral NP, i.e. the main IB component (22). The authors monitored the experimentally infected boas for two years, and in addition to IB formation in several tissues, the boas demonstrated virus secretion via feces and urates, but remained clinically healthy (22).

Based on the Schumacher and Wozniak studies (3, 4), and following anecdotal evidence of breeders employing pythons as sentinels of BIBD, since they can rapidly develop CNS signs, we initially made an attempt at experimental infection of juvenile pythons (*P. regius*). The experimental inoculation of the first set of pythons (N=8) revealed that inoculation via trachea or by intraperitoneal injection results in virus replication in multiple tissues. We considered inoculation via trachea to better mimic the natural infection route, and thus used tracheal instillation in the subsequent experiments. In the second experimental infection of pythons (*P. regius*), one individual developed severe CNS signs and we could re-isolate the virus from the brain of this snake, however, none of the snakes demonstrated IBs even after four months post inoculation. For the third experimental infection, 16 boa siblings (*B. constrictor*) were available and we co-housed two boas with pythons that had been experimentally infected prior to initiation of this experiment, however, we could not confirm horizontal transmission. Considering the results of Stenglein and co-workers, transmission from pythons to boas during co-housing can be considered as unlikely, since the authors did not find reptarenavirus RNA in python excreta (22). We immunized four boas prior to virus inoculations, and used a higher amount of virus for the tracheal inoculations. During the experiment, we learned that snakes with BIBD often harbor several reptarenaviruses, and that our “UHV inoculum” actually contained UHV-1 and ABV-1 at a roughly 1:1 ratio. On top of the co-infection with two distinct reptarenaviruses, we then superinfected some of the snakes with a genetically distinct reptarenavirus (UHV inoculated snakes re-inoculated with UGV-1) at approximately 3.5 months after the initial inoculation. We also attempted co-housing to demonstrate horizontal transmission, but none of the snakes developed IBs during the 10-month surveillance period, even though some boas tested RT-PCR positive for multiple reptarenaviruses. Our findings on pythons were similar to those of Stenglein and colleagues (22); however, unlike in other studies (3, 4, 22), we did not detect IB formation in the boas. On the other hand, our findings in the pythons, i.e. the clinical evidence that they are more prone to develop CNS signs than boas upon infection, concur with those made in earlier studies (3, 4, 22).

To understand why the boas did not develop IBs in our experimental set up, we studied the antibody response against NP, the main protein component of the IBs, and NAbs. We observed that the experimentally infected boas had mounted a strong antibody response against NP while lacking IBs. In our earlier study, we found low anti-NP antibody responses in snakes with BIBD (14), which together with the findings of the present study suggest that the antibody response could play a role in disease development. Interestingly, the amount of NAbs appeared to have a rather inverse correlation to the appearance of IBs, the experimentally infected snakes showed lower amounts of NAbs than the naturally infected snakes with the disease. Viremia in snakes with BIBD regardless of a strong NAbs response is interesting, and may be indicative of the role of snakes as reptarenavirus reservoirs, since persistently infected *Calomys musculinus* (dryland vesper mouse), the primary reservoir host of Junin virus (JUNV), a mammarenavirus, also possess NAbs (28-30). The same holds true for the persistently infected hantavirus rodent hosts (31, 32). The studies on JUNV and lymphocytic choriomeningitis virus (LCMV) suggest that mutations to the targets of NAbs could at least partially explain the persistence (28-30, 33). In our study, vaccination of snakes with either inactivated virus or recombinant NP resulted in strong antibody responses, however, the vaccination only protected one of the snakes against challenge with infectious virus. Unfortunately, our animal experimentation permits did not allow blood collection via cardiac venipuncture, due to which we obtained only a low amount of blood after completing the immunizations and could thus not analyze NAb titers. It is possible that immunization of snakes with detergent-inactivated reptarenaviruses did not induce a high enough NAb response to sustain virus challenge. Weakly neutralizing or non-neutralizing antibodies against reptarenavirus GPs could also boost the infection via antibody-dependent enhancement (ADE) i.e. by enabling the virus to enter Fc receptor expressing cells. ADE could allow infection by viruses bearing the GPs of different reptarenavirus species, e.g. antibodies against UHV-1 GPs could facilitate infection by virions with UGV-1 GPs. Further studies are needed to reveal whether a NAb response can be induced by e.g. recombinant reptarenavirus GPs, and whether the NAb response would actually protect the snakes from virus challenge. Vaccines against reptarenaviruses do currently not exist, though an effective vaccine might allow reptarenavirus eradication and enable BIBD-free snake collections or controlling the disease signs.

## ACKNOWLEDGEMENTS

This work was supported by the following grants: Jenny and Antti Wihuri Foundation (to JH), Academy of Finland (grant number 308613 and 314119 to JH). It also received a contribution from the Schweizerische Vereinigung für Wild-, Zoo-und Heimtiermedizin (SVWZH).

